# eMAGMA: An eQTL-informed method to identify risk genes using genome-wide association study summary statistics

**DOI:** 10.1101/854315

**Authors:** Zachary F Gerring, Angela Mina-Vargas, Eske M Derks

## Abstract

Identifying genes underlying genetic associations of complex disease is challenging because most common risk variants reside in non-protein coding regions of the genome and likely alter the expression of target genes by disrupting tissue and cell-type specific regulatory elements. To address this challenge, we developed a methodological framework, *eQTL-MAGMA* (*eMAGMA*), that converts SNP-level summary statistics into gene-level association statistics by assigning non-coding SNPs to their putative genes based on tissue-specific eQTL information. We compared eMAGMA to three eQTL informed gene-based approaches—S-PrediXcan, FUSION, and SMR—using simulated phenotype data. Phenotypes were simulated based on eQTL reference data using GCTA for all genes with at least one eQTL at chromosome 1 (651 genes). We performed 10 simulations per gene. The eQTL-*h*^2^ (i.e., the proportion of variation explained by the eQTLs was set at 1%, 2%, and 5%. We found eMAGMA outperforms other gene-based approaches across a range of simulated parameters (e.g. the number of identified causal genes). When applied to genome-wide association summary statistics for major depression, eMAGMA identified substantially more putative candidate causal genes compared to other eQTL-based approaches. By integrating tissue-specific eQTL information, these results show eMAGMA will help to identify novel candidate causal genes from genome-wide association summary statistics and thereby improve the understanding of the biological basis of complex disorders.

## Introduction

Genome-wide association studies (GWAS) have identified thousands of single nucleotide polymorphism (SNP) loci associated with disease risk [1]. However, the functional relevance of most SNP loci remains unknown, due in part to their position in non-protein coding regions of the genome [2]. Mapping trait-associated SNPs to their nearest gene often fails to identify the functional gene due to long-range regulatory effects on the expression of genes, known as expression quantitative trait loci (eQTLs). Furthermore, gene-based mapping methods that rely on arbitrary genomic windows to assign SNPs to genes, such as MAGMA [3], do not allow inferences on causal genes.

In recent years, several methods have been developed to integrate GWAS and gene expression information to improve our understanding of the functional mechanisms that underlie statistical genetic associations [4–6], known as a transcriptome-wide association study (TWAS). These methods are now widely used as secondary analyses using software packages such as FUSION [7], S-PrediXcan [6], and summary data-based Mendelian Randomisation Analysis (SMR) [5], and have identified novel genes and mechanisms underlying a range of diseases [2, 8]. Both TWAS and S-PrediXcan rely on a two-stage regression procedure. In the first stage, they train multi-variant prediction models in a sample with both genotype and gene expression data. In the second stage, these weights are then combined with summary-level data from GWAS to perform association analysis of estimated gene expression with a phenotype. SMR and its extension, the HEIDI test, aims to test for pleiotropic association between the expression level of a gene and a complex trait of interest using summary-level data from GWAS and expression quantitative trait loci (eQTL) studies within a Mendelian Randomization framework.

TWAS methods test the association between genetically determined component of gene expression and disease risk, ideally removing unwanted influences of environmental and technical factors on gene expression. However, this means only those genes whose expression can be reliably imputed from genotype data (i.e. moderately-highly heritable genes) can be tested for an association with a trait. Indeed, only 6,759 genes in GTEx (v7) whole blood—a relatively highly powered tissue—can be tested using S-PrediXcan, and 2,058 genes using FUSION. This drastically reduces the search space for prioritising candidate causal genes. We therefore created an alternative method, called eMAGMA, which modifies the MAGMA pipeline by mapping variants to genes based on tissue-specific eQTL information. We have used eQTL information from 48 tissues of the GTEx reference panel version 7 [9], although the method can be easily extended to other eQTL reference datasets. This approach was developed to identify functional gene associations that may be missed using proximity-based SNP assignment in MAGMA and may therefore identify alternative causal pathways from SNPs to trait.

Although a recent study performed a head-to-head comparison of two TWAS approaches, S-PrediXcan and FUSION [10], it was primarily based on observed GWAS data, and no systematic comparison was done to assess the performance of each method under different statistical parameters. We introduce the eMAGMA gene-based annotation approach and perform a systematic comparison of four different methods using data simulations and a real-life example using summary statistics from a GWAS of major depressive disorder. Our aims are to: (1) compare the statistical power of eMAGMA and other gene-based methods to detect a true association; (2) compare type-I error rates; (3) test the influence of the number of eQTLs on statistical power (i.e.weak instrument bias); and (4) compare gene-level effect sizes across methods. We plan to extend our simulations by modelling the performance of each method across different estimates of trait heritability and prevalence, and the proportion of overlap between causal GWAS variants and eQTL variants. A tutorial and input files are made available in a github repository: https://github.com/eskederks/eMAGMA-tutorial.

## Methods

### Gene-based methods

We compared four gene-based methods: S-PrediXcan [6], FUSION [7], SMR (version 1.0) [5], and our newly developed eMAGMA [11]. S-PrediXcan and FUSION are prediction-based approaches that impute the genetically regulated component of gene expression from SNP genotype data and regress the imputed expression on a given phenotype. SMR uses a Mendelian randomisation approach to estimate the effect of gene expression on a phenotype due to a single genetic marker (i.e. SNP), and tests whether a SNPs association with gene expression is due to linkage or pleiotropy (HEterogeneity In Depedent Instruments [HEIDI] test). MAGMA simply links SNPs to genes based on physical proximity, before combining the SNP-level P values while adjusting for linkage disequilibrium, gene size, and gene density. Our eMAGMA approach leverages significant (FDR<0.05) tissue-specific cis-eQTL information from GTEx (v7) to assign SNPs to putative genes.

### SNP genotype data for simulation analyses

The original genotype file from the QIMR Adult Twin Study [12, 13] included 3,738,240 SNPs from 28,110 individuals. We excluded non-founders (N=20,825), SNPs with > 1% missingness (N=1,023,785), and SNPs with minor allele frequency (MAF) < 0.05 (N=2,653,824). We subsequently excluded individuals with > 1% missing data (N=147). SNP identifiers were transformed to chr_chrposition to enable matching with GTEx eQTL reference data. This resulted in 43 duplicate SNPs, which were excluded from further analysis. Finally, we selected only SNPs from chromosome 1. The cleaned dataset included 7,138 subjects and 60,585 SNPs. eQTL information was obtained from whole blood samples of the GTEx eQTL database. (Whole_Blood.v7.signif_variant_gene_pairs.txt.gz). Significant eQTLs (FDR<0.05) were included in subsequent analyses. This eQTL refererence database included 655,939 eQTL-gene combinations for 8,235 unique genes.

### Phenotype simulation

Phenotypes were simulated using GCTA [15] using genotype and eQTL reference data from chromosome 1 (N=811 genes). For each gene, a phenotype was simulated using all significant (FDR<0.05) *cis*-eQTLs as predictors, based on the eQTL regression coefficients from the GTEx reference dataset. We performed 10 simulations per gene. Only those genes with at least one significant eQTL are included in the analysis (N=651). The eQTL-*h*^2^ (i.e., the proportion of variation explained by the eQTLs was set at 1%, 2%, or 5%.

### GWAS analysis

GWAS analyses of the 6,510 generated phenotypes were performed using the linear regression option in Plink [16]. SNP identifiers were replaced with rs identifiers using a lookup table to enable alignment with the annotation files in subsequent statistical analyses. We used the same significance level (*P*=6.25 × 10^−5^) for all analyses and corrected for the total number of genes in the GTEx whole blood reference dataset located at chromosome 1 (i.e. 0.05/811=1.2e-3).

### eMAGMA gene-level analysis

Since we are primarily interested in identifying functional variants associated with complex disorders, we leveraged eQTL data from 48 tissues in GTEx (version 7). Using the tissue-specific GTEx datasets, we generated SNP-gene pairs (FDR<0.05) that reflect functional relationships between SNPs and genes (cis-eQTLs), which serves as an input annotation file for the MAGMA software. We use the statistical framework from MAGMA to calculate gene-based P values using the updated (eQTL) annotation files. Gene-level analysis was done using default parameters and snp-wise=mean gene analysis model. We share comprehensive instructions on how to run eMAGMA in a github repository: https://github.com/AngelaMinaVargas/eMAGMA-tutorial

### Real-life example

We compared each method using observed GWAS summary statistics for major depressive disorder [14]. We assessed the correlation between the test statistics of each method using Pearson’s correlation coefficient.

### Comparative gene-level analyses

For the comparative analysis with S-PrediXcan, FUSION, and SMR, we applied prediction models trained in whole blood (GTEx v7) to analyse the generated simulated phenotype files, using the gene expression weight files provided by each package. We used software-specific default options for our analyses and used 1000 Genomes [17] data as the reference panel. We specified an annotation window 5kb upstream and 1.5kb downstream of each gene. For the eMAGMA annotation, we assigned SNPs to genes using significant (FDR<0.05) eQTL data from GTEx (v7). Gene-level analyses for eMAGMA was done using default parameters and snp-wise=mean gene analysis model.

## Results

We first counted the number of genes included in each post-GWAS method (Table 1). Interestingly, FUSION included the smallest number of genes, which is most likely due to the training algorithm excluding a large number of genes of which gene expression could not be imputed with sufficient accuracy. FUSION is therefore limited by the number of genes for which genetically-regulated gene expression can be reliably imputed.

**Table 1:** Number of genes and causal genes in each gene-based method.

| Method | Number of genes in model | Causal genes<br>(from simulations) | Proportion of causal genes |
| --- | --- | --- | --- |
| eMAGMA | 565 | 530 | 0.81 |
| MetaXcan | 628 | 490 | 0.75 |
| SMR | 401 | 387 | 0.59 |
| FUSION | 228 | 186 | 0.29 |

A total of 6,510 (651 genes with 10 simulations each) causal genes were used as input for phenotypic simulations. We first assessed the false positive rate (type-I error) of each method (i.e. under simulated conditions with no significant eQTLs/non-eQTLs) (Figure S1), and found all methods showed good control of the type-I error rate. We subsequently evaluated statistical power to detect association at a gene-based level, for varying levels of eQTL-h^2^. We assessed the proportion of significant associations relative to both the total number of causal genes (Figure 1) and when accounting for the total number of causal genes included in each method (Figure 2). eMAGMA outperformed all methods across different proportions of variance explained by the phenotype. After correcting for the number of genes included in each gene-based method, eMAGMA still outperformed other methods (Figure 2).

**Figure 1:**
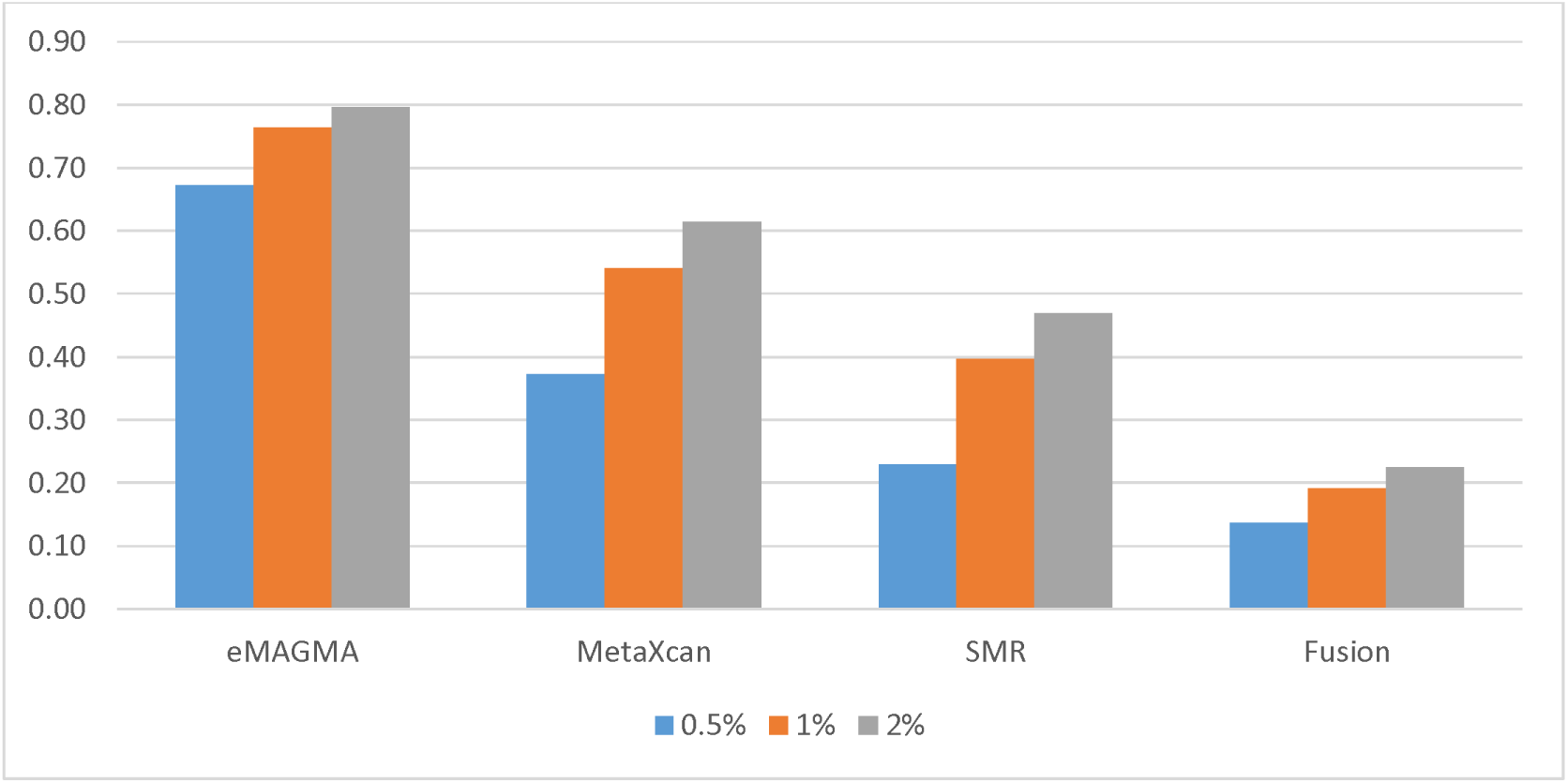
Proportion of significant associations (relative to the total number of causal genes)

**Figure 2:**
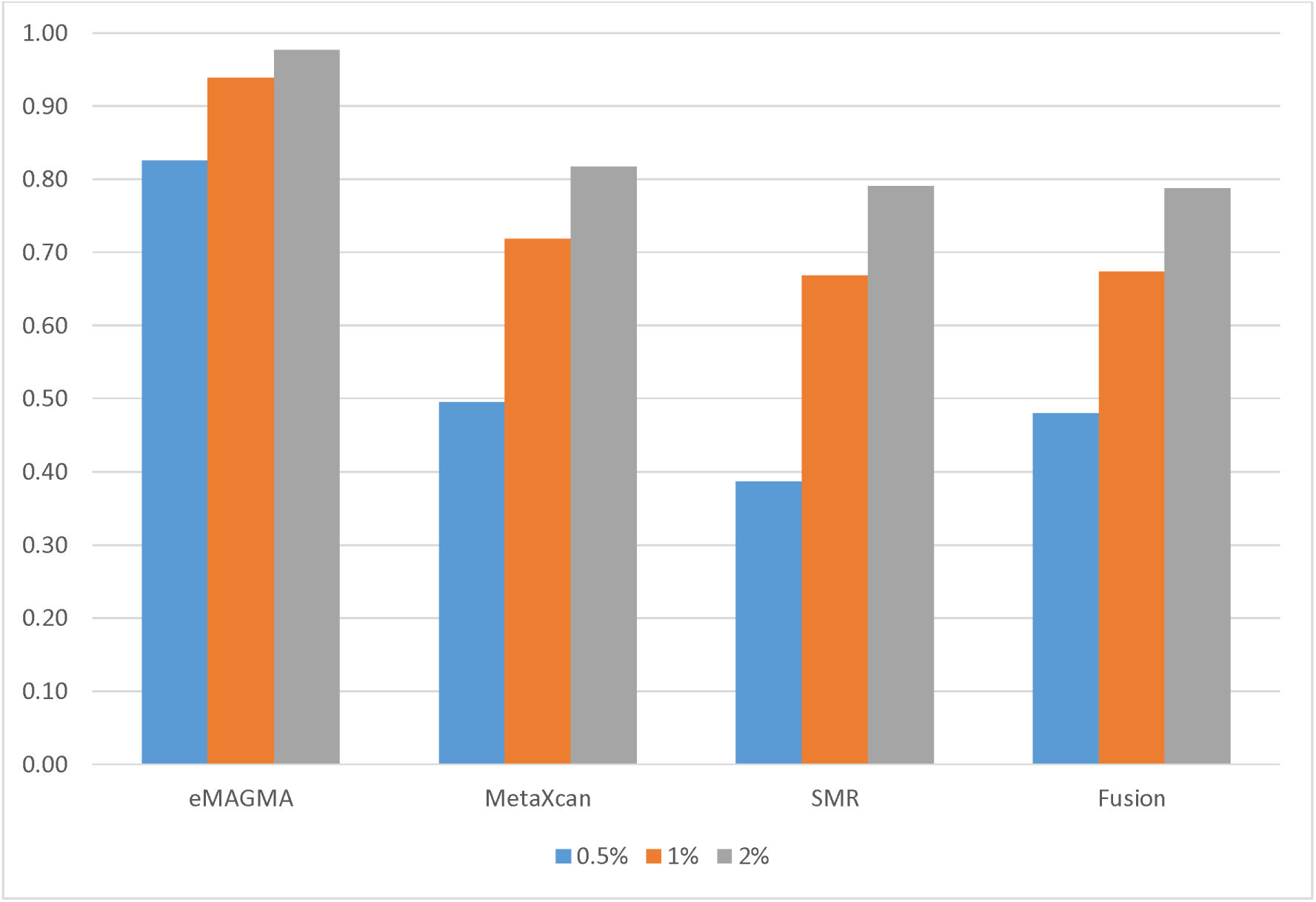
Proportion of significant associations (relative to the total number of causal genes per method)

All of the gene-based methods, with the exception of SMR, combine statistical evidence across multiple SNPs to derive a gene-based association. We therefore estimated statistical power as a function of the number of eQTLs per gene (Figure 3), with 1% of phenotypic variance explained by eQTLs. Power significantly increased with the number of eQTLs per gene (Figure 3; Supplementary Table S1). There was a significant association between the number of eQTLs per gene and statistical power for all methods, however eMAGMA was less sensitive to the number of eQTLs than the other methods.

**Figure 3:**
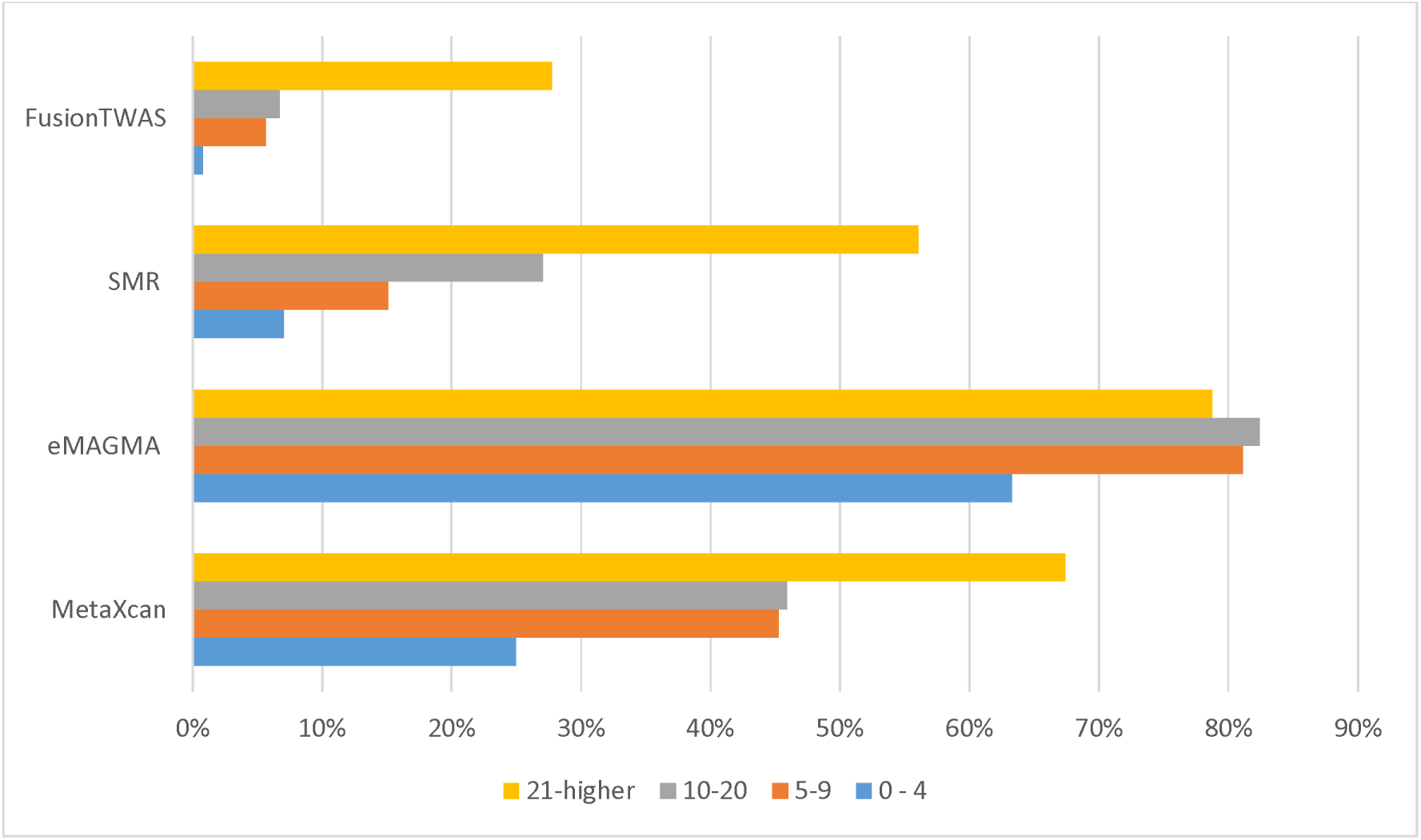
Statistical power as a function of the number of eQTLs per gene.

We assessed the overlap in genes between eQTL-based methods at 1% of phenotypic variance explained (Figure 4). The number of genes unique to each method far outweighed the overlap between any two methods, however there was good overlap across all four methods (n=851 from a total of 6,511 tests). We calculated the pairwise correlation of the Z-scores between gene-based methods (Table 2). Effect sizes of transcriptome-imputation methods were strongly correlated, particularly S-PrediXcan and FUSION (r=0.97, *P* < 2.2 × 10^−16^, df=168), but only low-moderate correlation was observed with eMAGMA (e.g. S-PrediXcan vs. eMAGMA; r=0.50, p<2.22 × 10^−16^, df=429) (Table 2).

**Table 2:** Correlation of Z-scores across 4 gene-based association methods.

|  | MetaXcan | eMAGMA | SMR | FUSION |
| --- | --- | --- | --- | --- |
| MetaXcan | 1 |  |  |  |
| eMAGMA | 0.50 | 1 |  |  |
| SMR | 0.78 | 0.31 | 1 |  |
| FUSION | 0.97 | 0.60 | 0.83 | 1 |

Notes: Correlation of Z-scores. Since eMAGMA Z-statistics do not reflect direction of effect, the absolute value of the Z-scores were used when correlating with eMAGMA results

**Figure 4:**
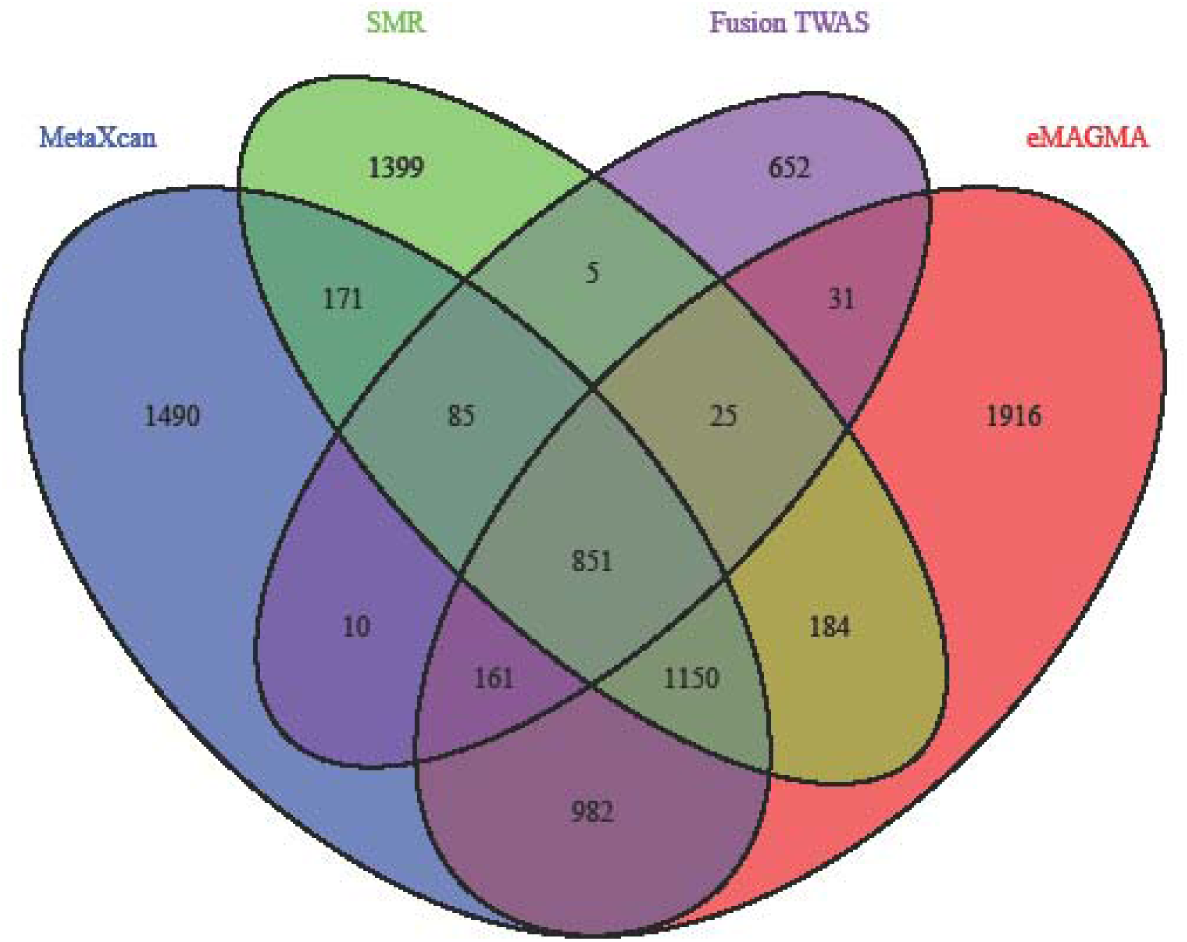
Overlap in genes between eQTL-based methods at 1% of phenotypic variance explained.

We compared the number of putative risk genes detected by each gene-based approach using GWAS summary statistics for Major Depression (Table 3).

**Table 3:** A real-life example comparing methods regarding the total number of significant associations with Major Depression.

| Method | Number of significant<br>Associations |
| --- | --- |
| eMAGMA | 85 |
| MetaXcan | 28 |
| SMR | 2 |
| FUSION | 14 |

## Discussion

We developed a gene-based method called eMAGMA which uses functional tissue-specific eQTL information from GTEx to assign SNPs to genes with the aim of improved annotation and interpretation of GWAS association signals. Our approach uses the statistical framework from MAGMA, but rather than assigning SNPs to genes based on physical proximity during gene annotation (i.e. mapping SNPs to genes using a pre-defined and arbitrary genomic window), we use significant (FDR<0.05) SNP-gene expression associations (eQTL) in GTEx. Our extension therefore provides more biologically meaningful and interpretable results compared to conventional MAGMA. We compared eMAGMA to three other eQTL-informed gene-based approaches (S-PrediXcan, FUSION, and SMR) using both simulated and observed GWAS data. We show that eMAGMA maintains appropriate control of the type-I error rate while outperforming other methods in detecting causal associations.

We used the methodological framework of MAGMA because it is one of the most widely used secondary analyses for the interpretation of GWAS results. Furthermore, the framework can be modified to include any type of annotation that maps SNPs to genes. For example, recent work to integrate chromatin interaction data from relevant tissues using the MAGMA framework increased power to identify putative risk genes and biological pathways for a range of neuropsychiatric traits [18]. With the availability of tissue-specific multi-omic (transcriptome, chromatin, Hi-C, DNA methylation) datasets through projects such as GTEx [19] and psychENCODE [20], it will be possible to link SNPs to target genes using the most functionally relevant data and improve the biological interpretation of GWAS results.

Recent gene-based methods integrate genetic and transcriptomic information to estimate the effect of genetically determined gene expression on phenotypic variation. No systematic comparison of the three most commonly-used methods—S-PrediXcan, FUSION, and SMR—has been done. However, a head-to-head comparison of S-PrediXcan and FUSION found the former approach captured known effects of genotype on expression more frequently than FUSION, although both methods tended to produce highly correlated results when applied to the same dataset [10]. We would like to note, however, that these results are limited to genes for which expression weights are available in both methods. We found FUSION tests far fewer genes than S-PrediXcan, suggesting S-PrediXcan is more appropriate for gene discovery.

We found all of the tested methods maintained control of the false positive under simulated conditions, where no single variant contributes to phenotypic variation. Under simulated conditions where 0.5%, 1%, or 2% of the phenotypic variation was explained by eQTLs (or non-eQTLs), S-PrediXcan captured more causal genes compared to SMR and FUSION which performed poorly. The performance of each method improved when measured against the actual total number of causal genes tested, correcting for the fact that some methods test fewer genes than others. eMAGMA was least influenced by the number of eQTLs of a gene, while all other methods tended to show a monotonic relationship with the number of eQTLs.

Our framework provides a more functionally valid gene-based test of association for GWAS compared to conventional MAGMA. However, it is prone to many of the same limitations of existing eQTL gene-based approaches. Notably, eMAGMA is not immune from the effects of linkage—where two or more variants in linkage disequilibrium independently affect gene expression and phenotypic variation—and pleiotropic SNP effects—where a single casual variant affects both gene expression and phenotypic variation. Our method (and other comparable eQTL methods) may therefore yield non-causal SNP-gene associations at nearby genes in the disease-associated region. The power of eMAGMA is limited by the sample size of the annotation eQTL dataset. This is especially problematic with brain tissue eQTL datasets, which tend to be underpowered given the inaccessibility of brain tissue. The meta-analysis of multiple independent brain eQTL datasets, performed by the psychENCODE consortium [20], will improve the power and interpretation of eMAGMA. Our simulations might favour eMAGMA over the other TWAS approaches because the eQTLs used in the annotation files were derived from the same reference eQTL dataset (GTEx) used to simulate gene expression. Future simulations using independent reference eQTL datasets will be required to confirm the better performance of eMAGMA. Finally, gene expression is highly cell-type specific [21]. The use of bulk tissue eQTL datasets may therefore reduce power to identify cell-type specific disease signals. The use of existing [21] and impending [20] single cell expression datasets may therefore improve the resolution of eQTL-based gene-mapping.

Future work will refine both the methodological framework of eMAGMA and the simulated data comparisons. First, our simulations were developed to compare statistical power of transcriptome imputation methods with MAGMA and eMAGMA. The simulations might be improved upon by modelling the impact of the proportion of causal eQTLs that contribute to phenotypic variation; that is, how do the methods perform under scenarios where only a subset of cis-eQTLs contribute of gene expression variation. Furthermore, we will assess the performance of each method across different estimates of trait heritability and prevalence. These additional analyses will provide a biologically valid and comprehensive assessment of model performance. Second, the tissue-specificity of eMAGMA may provide novel insights into biological mechanisms of disease, but at the cost of limited sample size— and statistical power—of tissue-specific eQTL datasets. Future work will annotate genes with eQTL from larger datasets blood-based eQTL datasets to improve gene discovery, before prioritising genes using tissue-specific results.

In conclusion, we present a modified MAGMA framework, eMAGMA that aggregates eQTL summary statistics into gene level association statistics for gene-level analyses. Using simulated data, we showed eMAGMA has greater power to detect causal associations compared to other popular gene-based approaches, while maintaining appropriate control of the type I error rate. Therefore, eMAGMA can provide a functionally relevant alternative to existing methods to identify genes and pathways from GWAS. A tutorial and input files can be found in the github repository: https://github.com/eskederks/eMAGMA-tutorial

